# Whole Genome Sequencing Analysis of a Recent Multi-Drug Resistant *Shigella sonnei* Outbreak Among Tunisian Children

**DOI:** 10.1101/2024.08.12.607540

**Authors:** Sana Ferjani, Lamia Kanzari, Zaineb Hamzaoui, Laila Jomni, Asma Ferjani, Ahmed Fakhfekh, Olfa Bahri, Wafa Achour, Mariem Zribi, Sameh Trabelsi, Nissaf Ben Alya, Dana Itani, Safa Bouwazra, Ebenezer Foster-Nyarko, Ilhem Boutiba Ben Boubaker

## Abstract

**Background:** *Shigella sonnei*, a leading cause of shigellosis, is a global health concern, particularly affecting children under five. The emergence of multidrug-resistant (MDR) strains, including resistance to key antibiotics like ciprofloxacin and third-generation cephalosporins, exacerbates treatment challenges. This study investigates the genetic and antimicrobial resistance profiles of *S. sonnei* isolates from Tunisia, focusing on an outbreak of extended-spectrum beta-lactamase (ESBL)-producing strains.

**Methods:** We analysed nine *S. sonnei* isolates collected between September 2022 and January 2023 from Tunisian hospitals, using whole genome sequencing (WGS). Standard bacterial identification and serotyping methods were employed alongside antimicrobial susceptibility testing. We examined the genetic relatedness of the isolates, identified resistance genes, and characterised virulence factors.

**Results:** All the isolates were confirmed as *S. sonnei* H6, biotype a, and belonged to lineage 3, clade 6 and sub-lineage 3. All harboured *bla*_CTX-M-15_, conferring resistance to third-generation cephalosporins. These were chromosomally integrated, suggesting stable resistance. Five isolates exhibited fluoroquinolone resistance associated with the *qn*rS1 gene, and all isolates had a single quinolone resistance-determining region mutation (GyrA-D87Y). Additionally, the plasmid-borne *mph*A gene, conferring resistance to macrolides, was prevalent. Single-linkage hierarchical clustering analysis indicated close genetic relationships with *S. sonnei* strains from Europe, particularly France and the UK (0 to 31 core genome MLST allele differences), indicating recent international dissemination.

**Conclusion:** This study provides the first comprehensive molecular characterisation of MDR *S. sonnei* in Tunisia, highlighting a significant public health threat. The findings underscore the importance of continuous genomic surveillance to track the spread of resistant strains and inform public health interventions.

## Introduction

*Shigella* spp. are human-adapted Gram-negative, non-spore-forming and non-motile, facultatively anaerobic pathogens that cause shigellosis—a severe gastrointestinal infection marked by profuse, often bloody diarrhoea accompanied by abdominal pain, fever, nausea, and vomiting (1). While known as a food- and travel-associated pathogen, in regions like Europe and North America, a substantial proportion of cases are sexually transmitted among men who have sex with men (2, 3). Globally, *Shigella* spp. accounts for significant morbidity and mortality, particularly affecting children under five years of age, with an estimated 75 million diarrhoeal episodes annually (4). The Global Enteric Multicentre Study found *Shigella* to be the most common cause of diarrhoea among children aged 2 to 5 years in sub-Saharan Africa and South Asia (5, 6).

The genus *Shigella* comprises four primary species, also known as subgroups or serogroups: *S. sonnei, S. flexneri, S. dysenteriae*, and *S. boydii*, each of which is further classified into serotypes and sub-serotypes based on antigenic differences in the somatic (O) of lipopolysaccharide expressed on the cell surface (1, 7). Notably, *S. sonnei* has become a dominant pathogen globally, prevalent in developed countries and the second most common in low- and middle-income countries, following *S. flexneri* (1, 8).

*Shigella spp*. exhibit a complex evolutionary history with three major clades (C1, C2, and C3) and *S. sonnei* as a notable outlier (8-10). *S. sonnei* is believed to have emerged more recently than other *Shigella* spp. and is more prevalent in developed countries, likely due to greater immunity in low-resource countries from exposure to *Plesiomonas shigelloides* O17, a faecal contaminant which shares an identical antigen (10).

An expansive study (11) utilising a diverse collection of *S. sonnei* isolates spanning more than six decades revealed several key insights into the population dynamics of this organism as follows. Based on phylogeographic analysis, *S. sonnei* appeared to have diverged from *Escherichia coli* around 1500 AD in central Europe and spread globally through travellers. Moreover, the population of *S. Sonnei* encompassed four lineages, Lineages 1 through 4. While the population from Europe spanned all four lineages, the bulk of the isolates from Africa, Asia and America fell into Lineage 3. Interestingly, Lineage 3 appeared to have emerged in Europe and spread to Africa in the early 1980s. Notably, the selection for multi-drug resistance (MDR) has been fundamental in driving this worldwide spread. Hawkey et al. (12) recently updated the lineages to include lineage 5 based on a comprehensive analysis of 1935 globally distributed genomes.

The emergence of MDR *S. sonnei* strains has raised significant concerns (2, 13). Historically, these strains have shown resistance to various antibiotics, including sulphonamides, ampicillin, streptomycin, and tetracycline, since the 1960s, conferred by an armamentarium of diverse antimicrobial resistance (AMR) genes (14). More recently, resistance to ciprofloxacin—the only first-line antimicrobial recommended by the World Health Organization (WHO) for empirical shigellosis treatment—alongside azithromycin and third-generation cephalosporins has been documented (14, 15). This evolution has led to the inclusion of these MDR strains on the WHO’s global priority pathogens list, highlighting an urgent need for novel antimicrobials (16, 17).

Resistance trends extend to extended-spectrum β-lactams, often linked to the acquisition of an extended-spectrum β-lactamase (ESBL) gene. This has been detected in sporadic cases and outbreaks across continents, including Australia, the UK, and Asia. An extensively drug-resistant (XDR) *S. sonnei* isolate, resistant to both the primary treatment ciprofloxacin and nearly all second-line antimicrobials, marked a significant outbreak in the UK in late 2021. Since then, numerous European countries have reported similar cases, predominantly among adult MSM. The transmission of MDR *S. sonnei* strains is facilitated by horizontal gene transfer mechanisms via resistance plasmids and other mobile genetic elements, as well as chromosomal pathogenic islands (18).

In Tunisia, shigellosis has occurred sporadically from 2000-2021, with an annual average of three cases. However, a notable increase in *S. sonnei* cases was reported in November 2022, with confirmed cases of ESBL-producing strains peaking in early January 2023, predominantly among children under five. This study utilises whole genome sequencing (WGS) to examine the *S. sonnei* strains isolated in Tunisia’s National Reference Laboratory of Antimicrobial Resistance Surveillance. The analysis aimed to determine the genetic relatedness of these isolates within both local and international contexts and identify their antimicrobial resistance markers and virulence factors.

## Methods

### Bacterial collection

Between September 2022 and January 2023, we observed a surge in reported cases of *S. sonnei* to Tunisia’s National Observatory of New and Emerging Diseases (19). By the first week of January 2023, 339 confirmed cases of ESBL-producing *S. sonnei* had been recorded nationwide. Nine isolates were submitted to the National Reference Laboratory for Antimicrobial Resistance Surveillance in Tunis for whole genome sequencing and further characterisation. These included seven isolates from the Charles Nicolle Hospital, one from the Aziza Othmana Hospital, and one from the National Bone Marrow Transplant Center, all in Tunis. Our analysis focused on these isolates. Additionally, we included a single quality control *S. sonnei* strain from 2010, isolated at the La Rabta Hospital in Tunis, to facilitate genomic comparisons with the outbreak strains (**Figure 1**).

**Figure 1:**
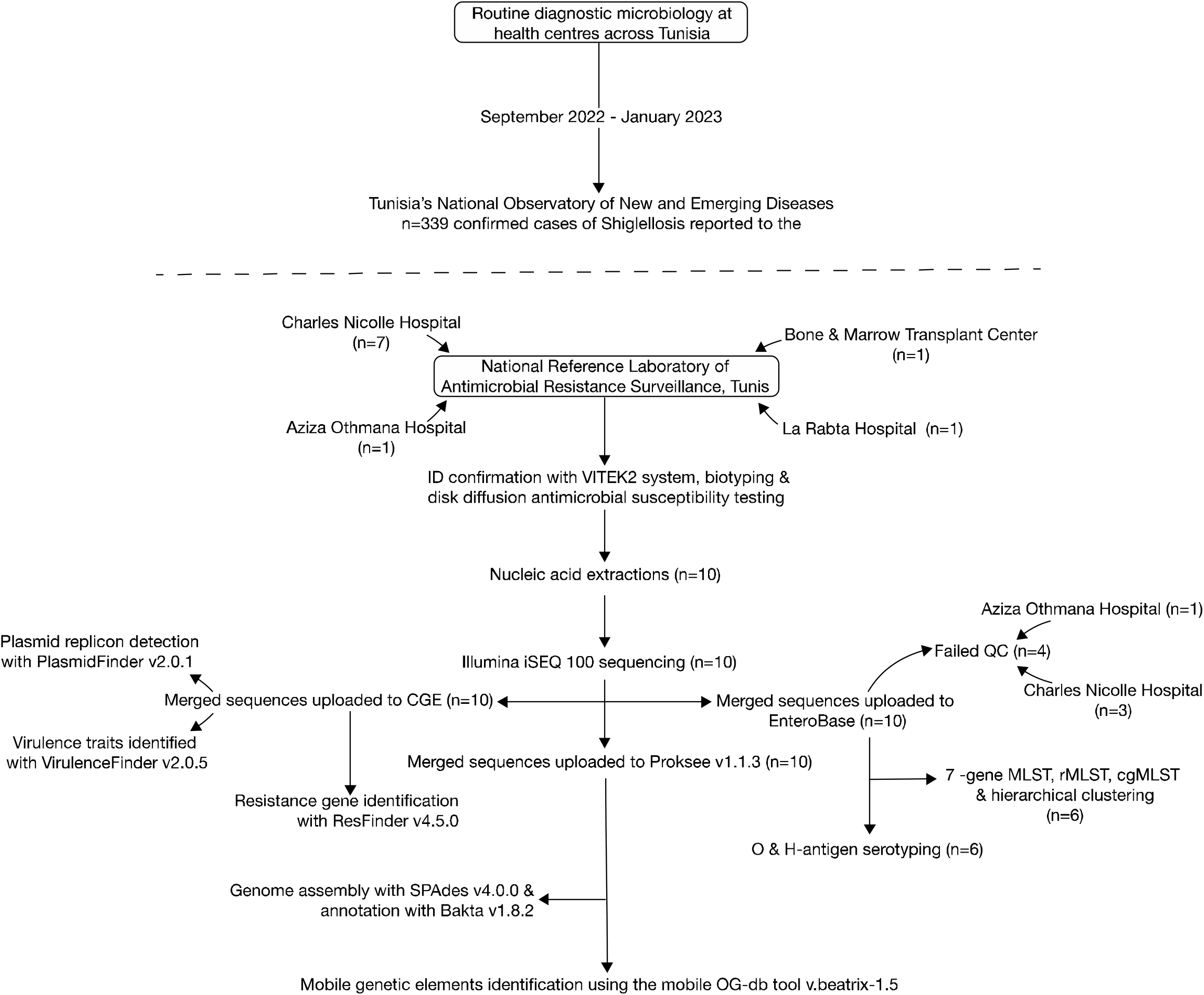
The study sample processing flow diagram.

All the isolates were derived from stool samples. Demographic and clinical data were retrieved from patient records, following approval from the respective department heads.

### Bacterial identification and antimicrobial susceptibility testing

Bacterial identification was conducted using conventional methods, including Gram staining, oxidase testing, and the VITEK2 system (BioMérieux, Marcy-l’Étoile, France). Agglutination tests were performed using *Shigella* polyvalent antisera (BioRad, Marnes-la-Coquette, France). Antimicrobial susceptibility profiles were determined via the standard disk diffusion method on Mueller Hinton agar, adhering to the EUCAST 2022 guidelines. The presence of extended-spectrum beta-lactamase (ESBL) production was screened using the double-disk synergy test.

### Sample processing for whole genome sequencing

#### DNA extraction

Each *S. sonnei* isolate was cultured in brain-heart infusion broth at 37°C for six hours. DNA was extracted from 1 ml of this culture using the Qiagen DNeasy Blood and Tissue Kit (Qiagen, Hilden, Germany), adhering to the manufacturer’s instructions. The DNA concentration and purity were assessed using a NanoDrop spectrophotometer.

#### Library preparation and sequencing

DNA was diluted to 5 ng/µL for library preparation, utilising the Illumina DNA Prep – (M) Tagmentation kit (Illumina). The concentration and quality of DNA libraries were measured using a Qubit fluorometer 2.0 and the QIAxcel system. Libraries were sequenced on the iSeq 100 System using the Reagent Kit v2 (300 cycles).

#### Sequencing data analysis

Raw sequencing reads were assembled *de novo* using Proksee Assemble software v1.3.0 via the Proksee v1.1.3 platform (20). Annotation of assembled genomes and plasmids was conducted using Bakta software v1.0.0 (21). The prevalence of mobile genetic elements was assessed using the mobile OG-db tool on the Proksee platform (20). The presence of antimicrobial resistance (AMR) and virulence genes were analysed using the ResFinder v4.5.0 (database version 2024-03-22) (22) and VirulenceFinder v2.0.5 (database version 2022-12-02) (23) web tools, while plasmid replicons were identified using PlasmidFinder v2.0.1 (database version 2023-01-18) (24), via the Center for Genomic Epidemiology integrated platform (https://www.genomicepidemiology.org/).

We utilised the hierarchical SNV-based genotyping scheme developed by Hawkey et al. (12), implemented via Mykrobe (25), to assign our isolates to the global *S. sonnei* lineages. For the determination of outer membrane (O) antigen and flagellin protein (H) types, Multilocus Sequence Types (MLST), ribosomal MLST (rMLST), and phylogenetic analysis to determine genetic neighbours based on core genome MLST (cgMLST), we uploaded the merged sequenced reads to EnteroBase integrative platform (26). For user-uploaded sequences, EnteroBase performs *de novo* assemblies using standardised pipelines (26), then assigned to stable clusters (HC0, HC2, HC5, up to HC2350) according to fixed cgMLT allelic distances using the single-linkage Hierarchical Clustering (HierCC) algorithm. Core genome MLST (cgMLST) profiling for *Escherichia coli* and *Shigella* utilises 2,512 core loci. Previous work has shown an 89% consistency between cgMLST allele differences among closely related genomes as determined by HierCC and core-genome single nucleotide polymorphisms (27). rMLST is based on 51-53 integers for ribosomal protein gene loci (28).

We searched for genomic relatives for each of our study isolates, beginning at HC0, through HC2, HC5 and upwards, until the closest relatives were identified. Next, we launched neighbour-joining trees for the identified genomic-relative clusters with NINJA – a hierarchical clustering tree-building algorithm that can scale to large (>100,000) input sequences (29).

To determine the lineages of our study isolates, we retrieved 196 genomes from the BioProject encompassing the strains used to delineate the global *S. sonnei* lineages (BioProject ID PRJEB2128), as defined by Holt et al. (11), along with 22 other publicly-available *S. sonnei* isolates from Tunisia. We then reconstructed a NINJA neighbour-joining tree to depict this collection and annotated the resulting tree in Adobe Illustrator 2024 v 28.0 (Adobe, San Jose, CA, USA).

## Results

### Epidemiologic and clinical data

All ten *S. sonnei* strains were isolated from children, predominantly females (n=8). Notably, six cases necessitated hospitalisation, presenting with symptoms ranging from diarrhoea with fever to acute functional renal failure and septic shock. A favourable progression was observed between three and ten days following appropriate antibiotherapy (**Table 1**).

**Table 1:**
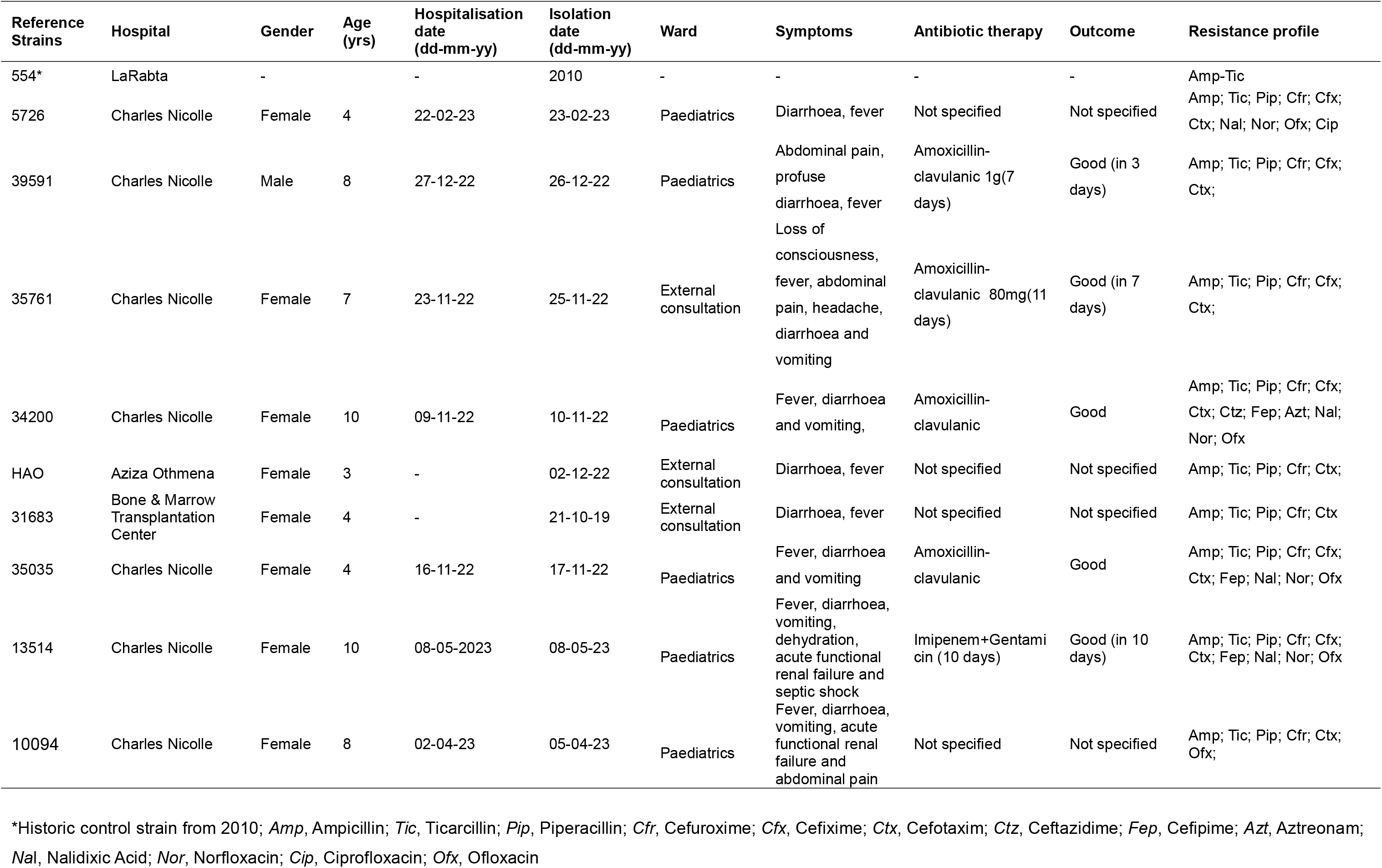
Epidemiological and clinical data of the Study *Shigella sonnei* isolates.

### Phenotypic characterisation

Biochemical identification using the VITEK2 system confirmed all strains as *S. sonnei*. Serotyping by agglutination identified all isolates as *S. sonnei* H6. Phenotypic biotyping revealed all nine outbreak strains as biotype a (ONPG+, rhamnose+, xylose−) and the single historic isolate as biotype g (ONPG+, rhamnose−, xylose−) (**Table S1**). All strains exhibited extended-spectrum beta-lactamase (ESBL) production and uniformly demonstrated phenotypic resistance to third-generation cephalosporins. Five isolates displayed phenotypic resistance to fluoroquinolones (**Table 1**).

### Plasmid incompatibility group and mobile genetic elements

Analysis of plasmid content identified the Col156 and IncFII plasmid types across all strains. ColRNAI and IncI1 were detected in 10094 and 35035 only. The distribution of genes associated with mobile genetic elements, such as integration/excision, replication/recombination/repair, phage, stability/transfer/defence, and transfer activities, varied, constituting between 9.9% and 16.4% of the total genomic content (**Table S2**).

### Prevalence of resistance genes

The resistance gene profile is detailed in **Table S2**. Apart from the historic isolate from 2010 (Isolate 554), all nine outbreak strains harboured *bla*_CTX-M-15_, along with *aph*(6)-Id, *dfr*A1, *mph*A, *qnr*S1; conferring resistance to third-generation cephalosporins, streptomycin, trimethoprim, macrolides, and quinolones, respectively. The *sul*2 and *tet*A genes, conferring resistance to sulfamethoxazole and tetracycline, respectively, were less prevalent, occurring in 4/10 (40%) and 5/10 (50%) of the study isolates, respectively.

The single chromosomal mutation, gyrA_D87Y, conferring reduced quinolone susceptibility, was found in all nine outbreak strains (**Table S2**). Genetic locations of *bla*_CTX-M-15_ and *qnr*S1 varied, with some colocalisation observed on chromosomes flanked by specific insertion sequences (**Figure S1**).

Additionally, all the study isolates uniformly encoded the *sit*A gene, conferring resistance to hydrogen peroxide.

### Virulence genes

A comprehensive analysis revealed 49 virulence genes coding for various functions, including toxins, adhesins, iron acquisition, and regulation of virulence expression (**Table S3**). All isolates contained the large virulence plasmid pINV, facilitating direct protein translocation into host cells. Notably, the *sig*A gene among the serine protease autotransporters of Enterobacteria (SPATEs) and genes critical for virulence regulation (*anr, nlp*I, *vir*B, *vir*F, *vir*K) were universally present. The distribution of invasion-associated genes was consistent across all isolates, except for the absence of *inv*E and *ial* genes.

### Clonality and phylogenetic analysis

All nine outbreak isolates were genotyped by Mykrobe as belonging to lineage 3, clade 6 and sub-lineage 3 (3.6.3), while the historic isolate typed as genotype 3.7.16.

Four of the ten study isolates (57260, 31683, HAO, and 10094) did not meet the EnteroBase quality metrics and were therefore excluded from the phylogenetic analysis. The remaining six isolates were identified as belonging to the ST152 clone and clustered with other lineage 3 genomes defined by Holt et al. (11) (**Figure 2**). Ribosomal MLST (rMLST) further classified these into rMLST 1458 and three novel rMLST types. Single-linkage hierarchical clustering identified the genetic relatives of our study isolates as follows.

**Figure 2:**
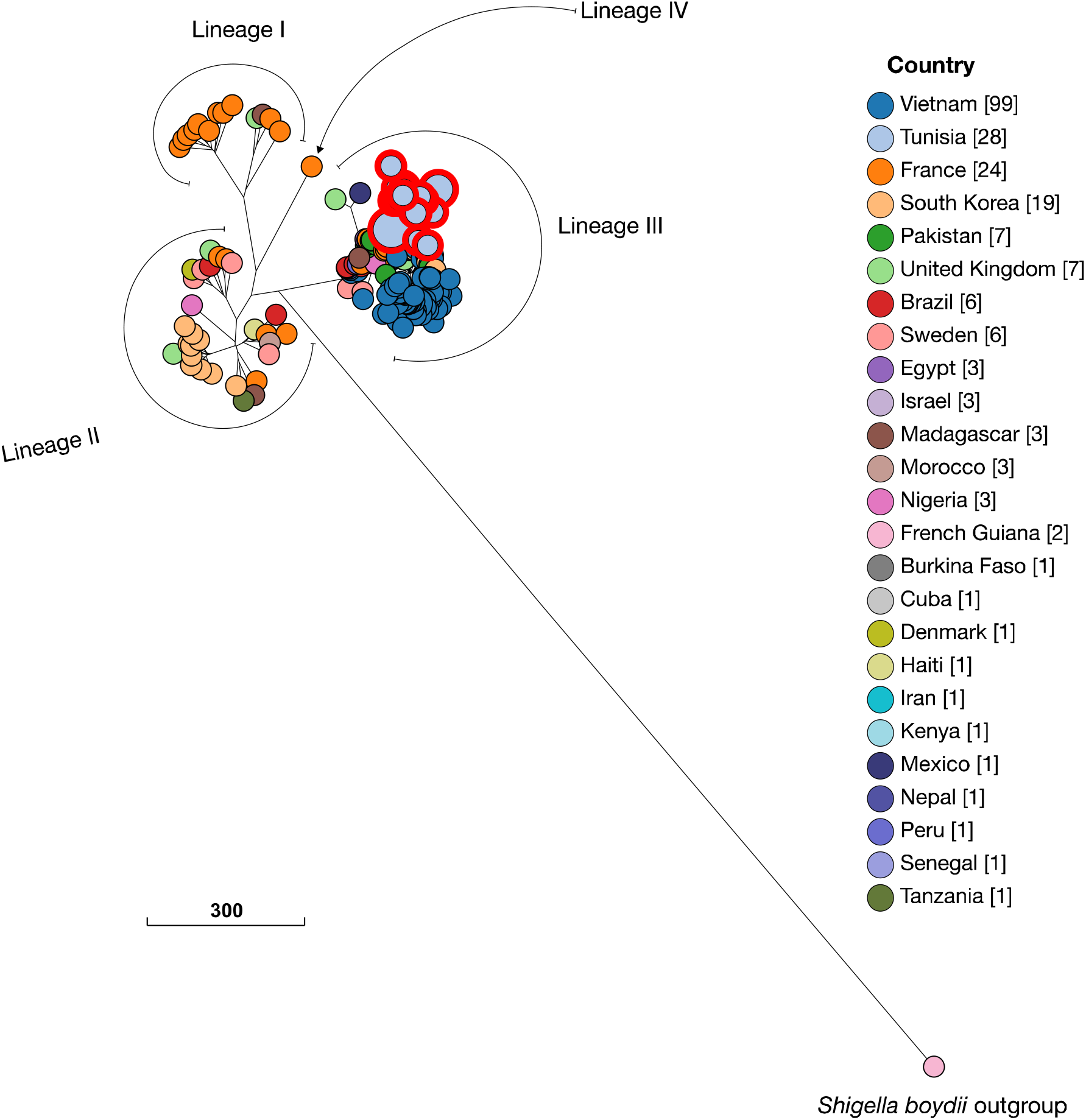
Global *Shigella sonnei* lineage classifications. The figure was generated using 196 global lineage markers as classified by Holt et al., (**Reference 11**), the six study isolates and 22 publicly available *S. sonnei* genomes from Tunisia collected between 2022 and 2023. The study isolates are highlighted in red and fall in Lineage III. The phylogenetic tree was reconstructed on the EnteroBase (**Reference 22**) integrative platform using the NINJA algorithm (Reference 28). The legend depicts the country of isolation, with the genome counts displayed in square brackets next to the country names.

Sample 13514 was indistinguishable from 3 genomes from France, isolated in 2023, and a single isolate from Chechia (isolated in 2022). These shared the same cgMLST type at level HC0, indicative of belonging to a recent common transmission cluster.

Isolate 35761 was equidistant from two close relatives: one from the UK and the other from France, separated by 6 alleles each. Similarly, isolate 39591 was separated from its nearest neighbours (one isolate from the UK and another from France) by ten alleles (**Figure 3A**).

**Figure 3:**
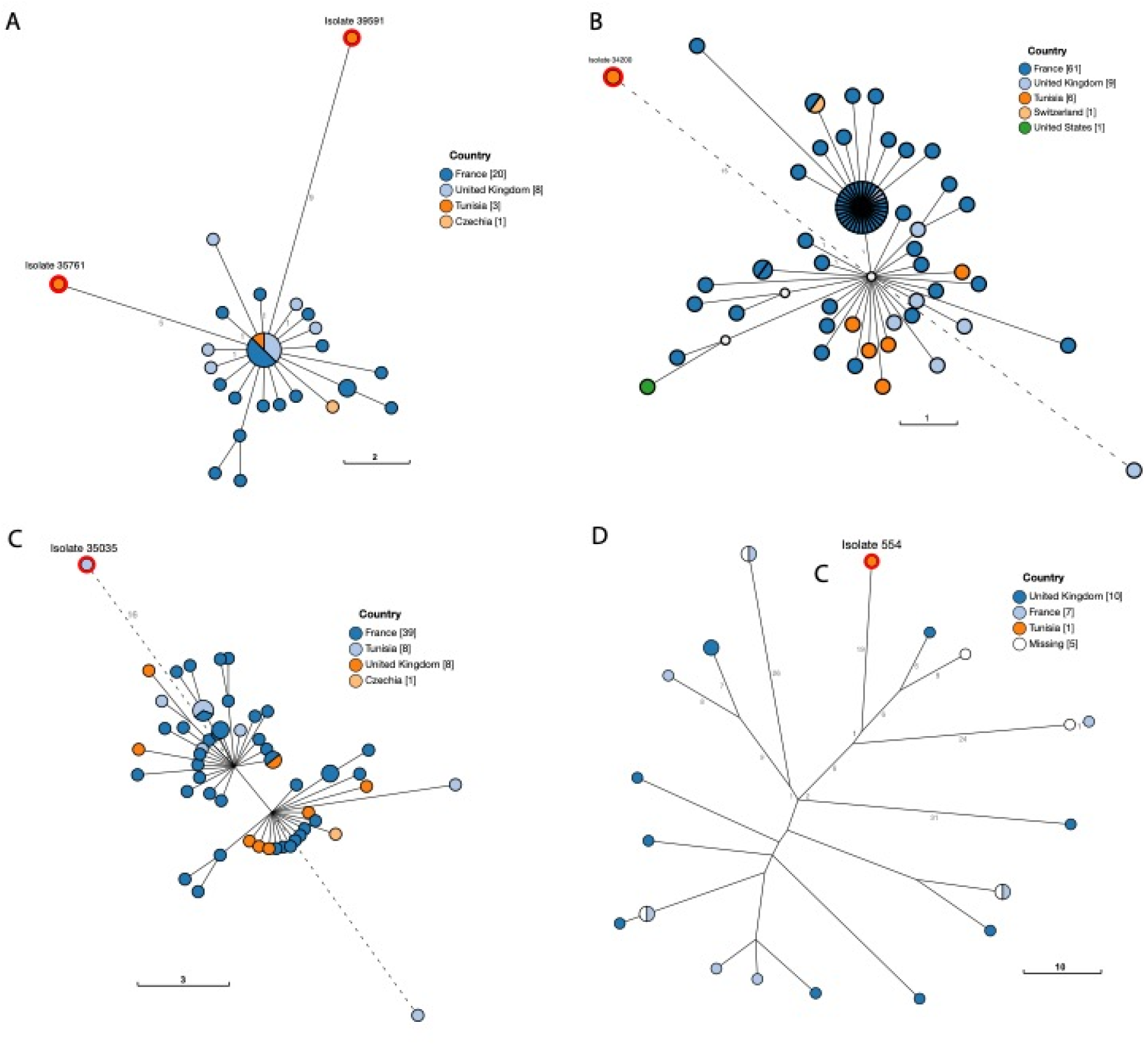
A NINJA neighbour-joining tree depicting the closest genomic relatives to the study isolates 35761, 39591, 34200, 35035, and 554. The study isolates are highlighted in red, with allelic distances to their closest relatives indicated by branch lengths. Study isolate 35761 is separated from its closest neighbour—an isolate from France—by 6 alleles. Isolate 39591 is 10 alleles away from its closest neighbours, one from the UK and another from France. Isolate 34200 is nested within isolates from France, separated from the closest neighbour by 16 alleles. Similarly, isolate 35035 is within a clade dominated by genomes from France, separated by 17 alleles from the nearest neighbour. Finally, isolate 554 is separated by 31 cgMLST alleles from its closest neighbour, an isolate from the UK. The legend shows the origin of isolation of the isolates, with the genome number presented in square brackets.

Isolate 34200 was likewise nested within isolates derived from France, with only 16 alleles separating it from its closest neighbour (**Figure 3B**). Isolate 35035 was separated from its closest neighbours by 17 cgMLST alleles, also isolated in France) in 2022 (**Figure 3C**). Finally, the historic isolate, 554, had its closest neighbour in a UK isolate, separated by 31 alleles (**Figure 4D**). These findings highlight a recent divergence and spread into Tunisia from Europe, in line with previous reports (11). Consistent with this, the publicly available Tunisian *S. sonnei* genomes consistently nested within clades dominated by genomes from France and the UK (0 to 5 cgMLST allele differences).

## Discussion

The protracted outbreak of ESBL-producing *Shigella sonnei* with a multidrug-resistant profile, including resistance to ciprofloxacin, poses a significant public health concern in Tunisia. This study marks the inaugural utilisation of whole-genome sequencing (WGS) at the national reference laboratory to investigate MDR *S. sonnei* isolates. WGS has demonstrated its crucial role as a precise tool for identifying *S. sonnei* strains, elucidating key epidemiological links, and identifying virulence and antimicrobial resistance genes.

Conventional identification and serotyping correlated perfectly with the molecular profiles of the strains, affirming the reliability of these methods, albeit within the confines of the limited number of isolates studied. Notably, all *S. sonnei* strains were ESBL producers, with a subset also exhibiting quinolone resistance. Genomic analyses revealed that these strains harboured *bla*_CTX-M-15_ in conjunction with genes encoding resistance to third-generation cephalosporins, streptomycin, trimethoprim, telithromycin, quinolones, and hydrogen peroxide.

The emergence of extended-spectrum β-lactamases (ESBLs) in *Shigella* species is a global concern, affecting both developed and developing regions (3, 30-33). A range of Ambler class A β-lactamases, including TEM, SHV, and CTX-M enzymes, has been reported among *Shigella* isolates (18). Our findings align with global trends, where *S. sonnei* frequently produces CTX-M-type enzymes, particularly CTX-M-15, documented in diverse geographic regions and associated with outbreaks. Notably, in our study, the *bla*_CTX-M-15_ gene was found chromosomally integrated, flanked by the insertion sequence IS26 and Tn2 transposase rather than located on plasmids. This chromosomal integration suggests a more stable resistance mechanism, as chromosomal genes are less likely to be lost than plasmid-borne genes. Additionally, this chromosomal location may reflect a historical integration event that has contributed to the persistence and dissemination of these resistance genes within our study population. This finding highlights the evolutionary adaptability of *S. sonnei* and underscores the necessity for continuous surveillance.

Ciprofloxacin, recommended as first-line treatment for dysentery by the WHO since 2005, has seen escalating resistance (13, 34-36). The first reported resistance in *S. sonnei* emerged around the turn of the millennium (18). Recent reports have indicated the acquisition of azithromycin resistance among ciprofloxacin-resistant *S. sonnei*, linked with high antibiotic use— representing significant challenges for empirical treatment of shigellosis (37, 38). Resistance is primarily due to mutations in the quinolone resistance determining regions (QRDR) of DNA gyrase and topoisomerase IV, although plasmid-mediated quinolone-resistance genes also play a role in enhancing resistance levels (18, 35, 36, 39, 40). Among our study isolates, five strains exhibited phenotypic fluoroquinolone resistance, with the *qnrS*1 gene detected in all strains, albeit chromosomally integrated, a factor likely to contribute to vertical, stable resistance transmission among clinically important lineages.

The aminoglycoside phosphotransferases encoded by *str*A and *str*B are prevalent among *Shigella* isolates and were observed in conjunction with *sul*2 and *tet*A genes on the identical plasmids in several of our study strains (41-44). Azithromycin, utilised as a second-line treatment in adults and a primary treatment in paediatric shigellosis, was compromised by the presence of the plasmid-mediated *mph*A gene, responsible for macrolide inactivation, in eight out of the ten isolates.

Phylogenetic comparisons using core genome multilocus sequence typing (cgMLST) with publicly available strains from Enterobase revealed that our isolates were closely related to those from Europe, particularly France and the UK. This finding highlights the potential for the international spread of *S. sonnei* clusters, likely facilitated by global travel, as documented in the literature (11). Consistent with previous studies, our isolates and other publicly available Tunisian strains clustered with the global *S. sonnei* lineage 3, suggesting the importation of this lineage from Europe to Tunisia (8, 11). However, in contrast to earlier reports indicating that biotype g predominates within lineage 3, most of our study isolates (nine out of ten) were classified as biotype a, with only one isolate belonging to biotype g. This discrepancy could be an artefact of our limited sample size, or it may indicate the presence of undersampled sub-population diversity within this globally dispersed lineage. Further research with larger sample sizes from our region is needed to explore this variation.

An obvious limitation of our study is the small sample size. Although many cases were detected across Tunisia during the study period, only a small fraction has been sequenced. Additionally, four samples could not be included in the phylogenetic analysis because they failed to meet the quality thresholds set by EnteroBase. Despite these challenges, we utilised the extensive collection of publicly available genomes on EnteroBase to enhance our analysis and provide a more comprehensive understanding of the *S. sonnei* population in our setting.

## Conclusion

Our investigation provides the first comprehensive molecular characterisation of MDR *S. sonnei* strains in Tunisia, highlighting the chromosomal integration of key resistance genes. Our findings underscore the critical need for ongoing epidemiological surveillance using WGS to monitor and manage the rising AMR burden in *S. sonnei*, a pathogen for which no vaccine is currently available. This study illustrates the local emergence of MDR strains and the potential for international dissemination, necessitating robust global cooperation in infectious disease monitoring and control.

## Data Summary

The raw sequences were uploaded to EnteroBase (https://enterobase.warwick.ac.uk/species/index/ecoli) and are available under assembly barcodes ESC_GB9141AA, ESC_GB9142AA, ESC_GB9146AA, ESC_GB9147AA, ESC_GB9148AA, a nd ESC_GB9149AA.

## Author notes

All supporting data, code, and protocols have been provided in the article or through supplementary data files. One supplementary figure and three supplementary tables are available in the online version of this article.

## Abbreviations

AMR: antimicrobial resistance
MDR: multi-drug resistance
MLST: multilocus sequence type
cgMLST: core genome multilocus sequence typing
WGS: whole genome sequencing
WHO: World Health Organization
ESBL: extended-spectrum β-lactamase gene
XDR: extremely drug resistant
QRDR: quinolone resistance determining region

## Impact Statement

*S. sonnei* is a major cause of dysentery prevalent in both developed and developing countries, with escalating resistance reported to essential antibiotics used for treatment, such as ciprofloxacin and azithromycin. In this study, we utilised whole genome sequencing to investigate a recent outbreak of multi-drug-resistant *S. sonnei* in Tunisia, revealing the potential international transmission and dissemination of the global *S. sonnei* lineage 3 of a lesser-known biotype, characterised by the chromosomal integration of resistance genes in the outbreak strains. Our research underscores the urgent need for ongoing genomic surveillance and targeted public health interventions to combat the spread of these resistant strains, particularly in paediatric populations, where treatment options are increasingly limited.

## Author contributions

Conceptualisation: IBBB, SF. Methodology: IBBB, SF, EFN. Formal analysis and visualisation: SF, LK, EFN. Data generation: OB, WA, MZ, ST, NBA, SB, LJ, AF, AF, DI. Data curation: IBBB, EFN. Supervision: IBBB. Funding: IBBB. Writing – Original draft preparation: SF, LK, EFN, IBBB. All authors read and approved the final manuscript.

## Ethical approval

The bacterial strains analysed in this study were isolated as part of routine diagnostic procedures; hence, ethical approval was not required. Informed patient consent was waived, as the samples were collected during standard diagnostic care. The research adhered to the principles outlined in the Helsinki Declaration.

## References

1. Kotloff KL, Riddle MS, Platts-Mills JA, Pavlinac P, Zaidi AKM. Shigellosis. Lancet. 2018;391(10122):801–12.

2. Mook P, McCormick J, Bains M, Cowley LA, Chattaway MA, Jenkins C, et al. ESBL-producing and macrolide-resistant Shigella sonnei infections among men who have sex with men, England, 2015. Emerg Infect Dis. 2016;22(11):1948–52.

3. Charles H, Prochazka M, Thorley K, Crewdson A, Greig DR, Jenkins C, et al. Outbreak of sexually transmitted, extensively drug-resistant Shigella sonnei in the UK, 2021-22: a descriptive epidemiological study. Lancet Infect Dis. 2022;22(10):1503–10.

4. Khalil IA, Troeger C, Blacker BF, Rao PC, Brown A, Atherly DE, et al. Morbidity and mortality due to Shigella and enterotoxigenic Escherichia coli diarrhoea: the Global Burden of Disease Study 1990-2016. Lancet Infect Dis. 2018;18(11):1229–40.

5. Kotloff KL, Nataro JP, Blackwelder WC, Nasrin D, Farag TH, Panchalingam S, et al. Burden and aetiology of diarrhoeal disease in infants and young children in developing countries (the Global Enteric Multicenter Study, GEMS): a prospective, case-control study. Lancet. 2013;382(9888):209–22.

6. Liu J, Platts-Mills JA, Juma J, Kabir F, Nkeze J, Okoi C, et al. Use of quantitative molecular diagnostic methods to identify causes of diarrhoea in children: a reanalysis of the GEMS case-control study. Lancet. 2016;388(10051):1291–301.

7. Wu Y, Lau HK, Lee T, Lau DK, Payne J. Serotyping based on whole-genome sequencing improves the accuracy of Shigella identification. Appl Environ Microbiol. 2019;85(7).

8. The HC, Thanh DP, Holt KE, Thomson NR, Baker S. The genomic signatures of Shigella evolution, adaptation and geographical spread. Nat Rev Microbiol. 2016;14(4):235–50.

9. Pupo GM, Lan R, Reeves PR. Multiple independent origins of Shigella clones of Escherichia coli and convergent evolution of many of their characteristics. Proc Natl Acad Sci U S A. 2000;97(19):10567–72.

10. Shepherd JG, Wang L, Reeves PR. Comparison of O-antigen gene clusters of Escherichia coli (Shigella) sonnei and Plesiomonas shigelloides O17: sonnei gained its current plasmid-borne O-antigen genes from P. shigelloides in a recent event. Infect Immun. 2000;68(10):6056–61.

11. Holt KE, Baker S, Weill FX, Holmes EC, Kitchen A, Yu J, et al. Shigella sonnei genome sequencing and phylogenetic analysis indicate recent global dissemination from Europe. Nat Genet. 2012;44(9):1056–9.

12. Hawkey J, Paranagama K, Baker KS, Bengtsson RJ, Weill FX, Thomson NR, et al. Global population structure and genotyping framework for genomic surveillance of the major dysentery pathogen, Shigella sonnei. Nat Commun. 2021;12(1):2684.

13. Mason LCE, Greig DR, Cowley LA, Partridge SR, Martinez E, Blackwell GA, et al. The evolution and international spread of extensively drug resistant Shigella sonnei. Nat Commun. 2023;14(1):1983.

14. Shad AA, Shad WA. Shigella sonnei: virulence and antibiotic resistance. Arch Microbiol. 2021;203(1):45–58.

15. Chung The H, Baker S. Out of Asia: the independent rise and global spread of fluoroquinolone-resistant Shigella. Microb Genom. 2018;4(4).

16. Tacconelli E, Carrara E, Savoldi A, Harbarth S, Mendelson M, Monnet DL, et al. Discovery, research, and development of new antibiotics: the WHO priority list of antibiotic-resistant bacteria and tuberculosis. Lancet Infect Dis. 2018;18(3):318–27

17. Tillotson G. A crucial list of pathogens. Lancet Infect Dis. 2018;18(3):234–6.

18. Ranjbar R, Farahani A. : Antibiotic-resistance mechanisms and new horizons for treatment. Infect Drug Resist. 2019;12:3137–67.

19. Tunisia’s National Observatory of New and Emerging Diseases. Monitoring and response bulletin. 1st August 2023. Available from https://onmne.tn/wp-content/uploads/2023/01/SHIGELLOSE-BULLTIN-N%C2%B01-2023.pdf.

20. Grant JR, Enns E, Marinier E, Mandal A, Herman EK, Chen CY, et al. Proksee: in-depth characterization and visualization of bacterial genomes. Nucleic Acids Res. 2023;51(W1):W484–W92.

21. Schwengers O, Jelonek L, Dieckmann MA, Beyvers S, Blom J, Goesmann A. Bakta: rapid and standardized annotation of bacterial genomes via alignment-free sequence identification. Microb Genom. 2021;7(11).

22. Zankari E, Hasman H, Cosentino S, Vestergaard M, Rasmussen S, Lund O, et al. Identification of acquired antimicrobial resistance genes. J Antimicrob Chemother. 2012;67(11):2640–4.

23. Liu B, Zheng D, Jin Q, Chen L, Yang J. VFDB 2019: a comparative pathogenomic platform with an interactive web interface. Nucleic Acids Res. 2019;47(D1):D687–D92.

24. Carattoli A, Zankari E, García-Fernández A, Voldby Larsen M, Lund O, Villa L, et al. In silico detection and typing of plasmids using PlasmidFinder and plasmid multilocus sequence typing. Antimicrob Agents Chemother. 2014;58(7):3895–903

25. Hunt M, Bradley P, Lapierre SG, Heys S, Thomsit M, Hall MB, et al. Antibiotic resistance prediction for Mycobacterium tuberculosis from genome sequence data with Mykrobe. Wellcome Open Res. 2019;4:191.

26. Zhou Z, Alikhan NF, Mohamed K, Fan Y, Achtman M, Group AS. The EnteroBase user’s guide, with case studies on Salmonella transmissions, Yersinia pestis phylogeny, and Escherichia core genomic diversity. Genome Res. 2020;30(1):138–52.

27. Frentrup M, Zhou Z, Steglich M, Meier-Kolthoff JP, Göker M, Riedel T, et al. A publicly accessible database for Clostridioides difficile genome sequences supports tracing of transmission chains and epidemics. Microb Genom. 2020;6(8).

28. Jolley KA, Bliss CM, Bennett JS, Bratcher HB, Brehony C, Colles FM, et al. Ribosomal multilocus sequence typing: universal characterization of bacteria from domain to strain. Microbiology (Reading). 2012;158(Pt 4):1005–15.

29. Wheeler TJ. Large-scale neighbor-joining with NINJA. in Algorithms in Bioinformatics. Berlin, Heidelberg: Springer Berlin Heidelberg. 2009.

30. Moreno-Mingorance A, Mir-Cros A, Goterris L, Rodriguez-Garrido V, Sulleiro E, Barberà MJ, et al. Increasing trend of antimicrobial resistance in Shigella associated with MSM transmission in Barcelona, 2020-21: outbreak of XRD Shigella sonnei and dissemination of ESBL-producing Shigella flexneri. J Antimicrob Chemother. 2023;78(4):975–82.

31. Campos-Madueno Edgar I, Bernasconi Odette J, Moser Aline I, Keller Peter M, Luzzaro F, Maffioli C, et al. Rapid increase of CTX-M-producing Shigella sonnei isolates in Switzerland due to spread of common plasmids and international clones. J Antimicrob Chemother. 2020;64(10):10.1128/aac.01057-20.

32. ECDC. European Centre for Disease Prevention and Control. Increase in extensively-drug resistant Shigella sonnei infections in men who have sex with men in the EU/EEA and the UK – 23 February 2022. ECDC: Stockholm; 2022.

33. Puzari M, Sharma M, Chetia P. Emergence of antibiotic resistant Shigella species: A matter of concern. J Infect Public Health. 2018;11(4):451–4.

34. World Health O. Guidelines for the control of shigellosis, including epidemics due to Shigella dysenteriae type 1. Geneva: World Health Organization; 2005.

35. Tajbakhsh M, García Migura L, Rahbar M, Svendsen CA, Mohammadzadeh M, Zali MR, et al. Antimicrobial-resistant Shigella infections from Iran: an overlooked problem? J Antimicrob Chemother. 2012;67(5):1128–33.

36. Tamanna Ramana, J. Structural Insights into the fluoroquinolone resistance mechanism of Shigella flexneri DNA gyrase and topoisomerase IV. Microb Drug Resist. 2016;22(5):404–11.

37. Malaka De Silva P, Stenhouse GE, Blackwell GA, Bengtsson RJ, Jenkins C, Hall JPJ, et al. A tale of two plasmids: contributions of plasmid associated phenotypes to epidemiological success among Shigella. Proc Biol Sci. 2022;289(1980):20220581.

38. Baker KS, Dallman TJ, Field N, Childs T, Mitchell H, Day M, et al. Horizontal antimicrobial resistance transfer drives epidemics of multiple Shigella species. Nat Commun. 2018;9(1):1462.

39. Chung The H, Rabaa MA, Pham Thanh D, De Lappe N, Cormican M, Valcanis M, et al. South Asia as a reservoir for the global spread of ciprofloxacin-resistant Shigella sonnei: A cross-sectional study. PLoS Med. 2016;13(8):e1002055.

40. Dutta S, Pazhani GP, Nataro JP, Ramamurthy T. Heterogenic virulence in a diarrheagenic Escherichia coli: evidence for an EPEC expressing heat-labile toxin of ETEC. Int J Med Microbiol. 2015;305(1):47–54.

41. Shaw PC, Liang AC, Kam KM, Ling JM. Presence of strA-strB gene within a streptomycin-resistance operon in a clinical isolate of Shigella flexneri. Pathology. 1996;28(4):356–8.

42. Sundin GW. Distinct recent lineages of the strA-strB streptomycin-resistance genes in clinical and environmental bacteria. Curr Microbiol. 2002;45(1):63–9.

43. Muthuirulandi Sethuvel DP, Anandan S, Devanga Ragupathi NK, Gajendiran R, Kuroda M, Shibayama K, et al. IncFII plasmid carrying antimicrobial resistance genes in Shigella flexneri: Vehicle for dissemination. J Glob Antimicrob Resist. 2019;16:215–9.

44. Parajuli P, Rajput MI, Verma NK. Plasmids of Shigella flexneri serotype 1c strain Y394 provide advantages to bacteria in the host. BMC Microbiol. 2019;19(1):86.

