## Supplementary material for "Whole Genome Sequencing Analysis of a Recent Multi-Drug Resistant *Shigella sonnei* Outbreak Among Tunisian Children": Figure S1

**Supplementary Figures**
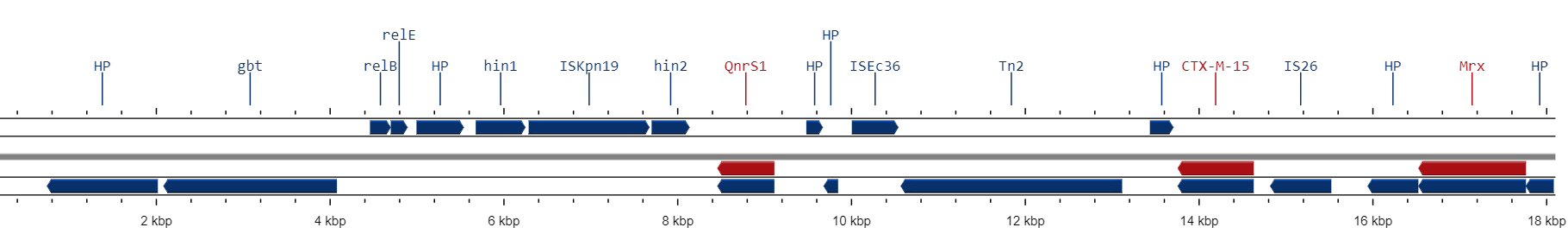

**Figure S1.** Map of the chromosomal DNA fragment carrying *bla*_CTX-M-15_ and *qnr*S1 resistance genes of *Shigella sonnei* strains. The figure was generated using the mobile OG-db tool on the Proksee platform (**Reference 20**). HP, hypothetical protein; GPT, Glycine betaine transporter; ISKpn19, ISKra4 family transposase; ISEc36, IS3 family transposase; IS26, IS6 family transposase; Tn2, Tn3 family transposase; Mrx is part of the macrolide inactivation gene.
