## Supplementary material for "Whole Genome Sequencing Analysis of a Recent Multi-Drug Resistant *Shigella sonnei* Outbreak Among Tunisian Children": Table S1; Table S2; Table S3

**Table S1:** Biotypes of the Study isolates (VITEK2 results)

|  | **554*** | **57260** | **39591** | **35761** | **35035** | **HAO** | **31683** | **34200** | **13514** | **10094** |
| --- | --- | --- | --- | --- | --- | --- | --- | --- | --- | --- |
| APPA | - |  |  |  |  |  |  |  |  |  |
| H2S | - | - | - | - | - | - | - | - | - | - |
| BGLU | - | - | - | - | - | - | - | - | + | - |
| ProA | + | - | - | - | - | + | + | - | - | + |
| SAC | - | - | - | - | - | - | - | - | - | - |
| ILATK | - | - | - | - | - | - | - | - | + | + |
| GlyA | - | - | - | - | - | - | - | - | + | - |
| O129R | + | + | + | + | + | + | + | + | + | + |
| ADO | - | - | - | - | - | - | - | - | - | - |
| BNAG | - | - | - | - | - | - | - | - | - | - |
| dMAL | + | - | - | - | - | - | + | - | - | + |
| LIP | - | - | - | - | - | - | - | - | - | - |
| dTAG | - | - | - | - | - | - | - | - | - | - |
| AGLU | - | - | - | - | - | - | - | - | - | - |
| ODC | + | - | - | - | - | + | + | - | + | + |
| GGAA | - | - | - | - | - | - | - | - | + | - |
| PyrA | - | - | - | - | - | - | - | - | - | - |
| AGLTp | - | - | - | - | - | - | - | - | - | + |
| dMAN | - | + | - | - | - | + | + | + | + | + |
| PLE | - | - | - | - | - | - | - | - | - | - |
| dTRE | + | + | + | + | + | + | + | + | + | + |
| SUCT | + | - | - | - | - | + | + | - | + | + |
| LDC | - | - | - | - | - | - | - | - | - | - |
| IMLTa | - | - | - | - | - | - | - | - | - | - |
| IARL | - | - | - | - | - | - | - | - | - | - |
| dGLU | + | + | + | + | + | + | + | + | + | + |
| dMNE | + | + | - | - | - | + | + | + | + | + |
| TyrA | + | - | - | - | - | + | + | + | + | + |
| CIT | - | - | - | - | - | - | - | - | - | - |
| NAGA | - | - | - | - | - | - | - | - | - | + |
| IHISa | - | - | - | - | - | - | - | - | - | - |
| ELLM | - | - | - | - | - | - | + | - | - | - |
| dCEL | - | - | - | - | - | - | - | - | - | - |
| GGT | - | - | - | - | - | - | - | - | - | - |
| BXYL | - | - | - | - | - | - | - | - | - | - |
| URE | - | - | - | - | - | - | - | - | - | - |
| MNT | - | - | - | - | - | - | - | - | - | - |
| AGAL | + | - | - | - | - | + | + | - | + | + |
| CMT | + | - | - | + | - | + | + | + | + | + |
| ILATa | - | - | - | - | - | - | - | - | - | - |
| BGAL | - | - | - | - | - | + | - | + | + | - |
| OFF | + | - | - | + | - | + | + | + | + | + |
| BAIap | - | - | - | - | - | - | - | - | - | - |
| dSOR | - | - | - | - | - | - | - | - | - | - |
| 5KG | + | - | - | - | - | + | + | - | + | + |
| PHOS | + | - | - | - | - | - | + | - | - | + |
| BGUR | + | - | - | - | - | + | + | - | + | + |
| Biotype | a | a | a | a | a | a | g | a | a | a |

*APPA*, Ala-Phe-Pro-Arylamidase; *H_2_S*, H_2_S production; *BGLU*, Beta-glucosidase; *ProA*, L-Proline Arylamidase; *SAC*, Saccharose/Sucrose; *ILATK*, L-Lactate alkalinization; *GlyA*, Glycine Arylamidase; *O129R*, O/129 Resistance; *ADO*, Adonitol; *BNAG*, Beta-N-Acetyl-Glucosaminidase; *dMAL*, D-Maltose; *LIP*, Lipase; d*TAG*, D-Tagatose; *AGLU*, ALPHA-Glucosidase; *ODC*, Ornithine Decarboxylase; *GGAA*, Glu-Gly-Arg-Arylamidase; *PyrA*, L-Pyrrolydonyl-Arylamidase; *AGLTp*, Glutamyl ArylamidasepNA; dMAN, D-Mannitol; *PLE*, Palatinose; *dTRE*, D-Trehalose; *SUCT*, Succinate alkalinization; *LDC*, Lysine Decarboxylase; *IMLTa*, L-Malate assimilation; *IAR*L, L-Arabitol; *dGLU*, D-Glucose; *dMNE*, D-Mannose; *TyrA*, Tyrosine Arylamidase; *CIT*, Citrate, Sodium; *NAGA*, Beta-N-Acetyl-Galactosaminidase; *IHISa*, L-Histidine assimilation; *ELLM*, Ellman; *dCEL*, D-Cellobiose; *GGT*, Gamma-Glutamyl-Transferase; *BXYL*, Beta-Xylosidase; *URE*, Urease; *MNT*, Malonate; *AGAL*, Alpha-Galactosidase; *CMT*, Coumarate; *ILATa*, L-Lactate assimilation; *BGAL*, Beta-Galactosidase; *OFF*, Fermentation/Glucose; *BAIa*p, Beta-Alanine Arylamidase; *dSOR*, D-Sorbitol; *5KG*, 5-Keto-D-Gluconate; *PHOS*, Phosphatase; *BGUR*, Beta-Glucuronidase; *Historic control strain from 2010

**Table S2:** Distribution of AMR genes, mobile genetic elements and plasmid typing profiles of the Study isolates

|  | **554*** | **57260** | **39591** | **35761** | **35035** | **HAO** | **31683** | **34200** | **13514** | **10094** |
| --- | --- | --- | --- | --- | --- | --- | --- | --- | --- | --- |
| AMR genes | | | | | | | | | | |
| *aad*A1 | + | - | - | - | - | - | - | - | - | - |
| *aph*(3'')-Ib | - | + | - | - | - | - | - | - | - | + |
| *aph*(6)-Id | - | + | + | + | + | + | + | - | - | + |
| *bla*_CTX-M-15_ | - | + | + | + | + | + | + | + | + | + |
| *dfr*A1 | + | + | + | + | + | + | + | + | + | + |
| *mph*(A) | - | + | + | + | + | + | + | - | + | + |
| *qnr*S1 | - | + | + | + | + | + | + | - | + | + |
| *sit*ABCD | + | + | + | + | + | + | + | - | + | + |
| *sul*2 | + | - | - | - | - | - | + | + | + | - |
| *tet*(A) | + | - | + | + | - | + | - | + | - | - |
| D87Y*gyrA* mutation | - | + | + | + | + | + | + | + | + | + |
| Plasmid typing | | | | | | | | | | |
| Col(BS512) | + | - | - | - | - | - | - | - | - | - |
| Col(MG828) | + | - | - | - | - | - | - | - | - | - |
| Col156 | + | + | + | + | + | + | + | + | + | + |
| IncFII | - | + | + | + | + | + | + | + | + | + |
| IncI1-I(Alpha) | - | - | - | - | + | - | - | - | - | - |
| ColRNAI | - | - | - | - | - | - | - | - | - | + |
| Mobile genetic elements (number of genes detected) | | | | | | | | | | |
| Integration/excision | 81 | 150 | 147 | 138 | 162 | 141 | 110 | 157 | 139 | 109 |
| Replication/recombination/repair | 125 | 141 | 137 | 138 | 162 | 146 | 133 | 145 | 138 | 139 |
| Phage | 126 | 126 | 118 | 118 | 153 | 140 | 127 | 122 | 118 | 108 |
| Stability/transfer/defense | 59 | 72 | 75 | 71 | 83 | 75 | 56 | 76 | 79 | 62 |
| Transfer | 58 | 96 | 98 | 92 | 179 | 101 | 60 | 98 | 95 | 83 |
| Total | 449 | 585 | 575 | 557 | 739 | 603 | 486 | 598 | 569 | 501 |

*aad*A1, *aph*(3'')-Ib, *aph*(6)-Id, streptomycin resistance; *bla*_CTX-M-15_, 3^rd^ generation cephalosporin resistance; *dfr*A1, trimethoprim resistance; *mph*(A), macrolide resistance; *qnr*S1, quinolones resistance; *sit*A/B/C/D, hydrogen peroxide resistance; *sul*2, sulfamethoxazole resistance; *tet*A, tetracycline resistance; * Historic control strain from 2010

**Table S3:** Distribution of virulence genes among the Study isolates

|  | | 554* | 57260 | 39591 | 35761 | 35035 | HAO | 31683 | 34200 | 13514 | 10094 |
| --- | --- | --- | --- | --- | --- | --- | --- | --- | --- | --- | --- |
| Diverse | *anr* | + | + | + | + | + | + | + | + | + | + |
|  | *cap*U | - | + | + | + | + | + | + | + | + | + |
|  | colE2 | + | - | - | - | - | - | - | - | - | - |
|  | colE7 | - | + | + | + | + | + | + | + | + | + |
|  | *gad* | + | + | + | + | + | + | + | + | + | + |
|  | *nlp*I | + | + | + | + | + | + | + | + | + | + |
|  | *sepA* | - | - | - | - | - | - | - | - | - | - |
|  | *ter*C | + | + | + | + | + | + | + | + | + | + |
|  | *tra*T | - | + | + | + | + | + | + | + | + | + |
|  | *vir*K | - | + | + | + | + | + | + | + | + | + |
|  | *vir*B | - | + | + | + | + | + | - | - | + | + |
| Invasion | *hly*E | + | + | + | + | + | + | + | + | + | + |
|  | *invE* | - | - | - | - | - | - | - | - | - | - |
|  | *ial* | - | - | - | - | - | - | - | - | - | - |
|  | *gtr* | - | - | - | - | - | - | - | - | - | - |
|  | *sen*B | + | + | + | + | + | + | + | + | + | + |
|  | *shi*A | + | + | + | + | + | + | + | + | + | + |
|  | *shi*B | + | + | + | + | + | + | + | + | + | + |
|  | *sig*A | + | - | + | + | + | - | + | + | + | + |
|  | *stx* | - | - | - | - | - | - | - | - | - | - |
|  | *vir*F | - | + | + | + | + | + | + | + | + | + |
|  | *vir*A | - | + | + | + | + | - | - | + | - | + |
|  | *Ipa* operon: Type III secretion system | | | | | | | | | | |
|  | *ipa*A | - | + | + | + | + | + | - | + | + | + |
|  | *ipa*B | - | + | - | + | + | + | - | + | + | + |
|  | *ipa*C | - | + | + | + | + | + | - | + | + | + |
|  | *ipa*D | - | + | + | + | + | + | + | + | + | + |
|  | *ipa*J | - | + | + | + | - | + | - | - | + | + |
|  | *ipa*H9.8/5 | -/+ | +/- | +/- | +/- | +/- | +/- | +/- | -/+ | +/- | +/- |
|  | *Osp* operon: Type III secretion system | | | | | | | | | | |
|  | *osp*A | - | - | - | - | - | - | - | - | - | - |
|  | *osp*B | - | + | + | + | + | - | - | + | + | + |
|  | *osp*C1 | - | + | + | + | + | - | + | - | - | + |
|  | *osp*D1/3 | - | +/+ | +/+ | +/- | +/+ | - | +/- | +/+ | +/- | +/+ |
|  | *osp*F | - | + | + | + | + | - | + | + | + | + |
|  | *osp*G | - | + | + | + | + | - | + | + | + | + |
|  | *osp*I | - | + | - | + | + | - | + | + | + | + |
|  | *osp*Z | - | + | - | + | + | - | + | + | + | + |
|  | *Ipg* operon: Type III secretion system | | | | | | | | | | |
|  | *ipg*A | - | + | - | + | + | + | - | + | + | + |
|  | *ipg*B1 | - | + | + | + | + | + | - | - | + | + |
|  | *ipg*C | - | + | + | + | + | + | - | + | + | + |
|  | *ipg*D | - | + | + | + | + | + | - | + | + | + |
|  | *ipg*E | - | + | + | + | + | + | - | + | + | + |
|  | *ipg*F | - | + | - | + | + | + | - | + | + | + |
|  | *ics* operon: Type III secretion system | | | | | | | | | | |
|  | *icsB* | - | + | - | + | - | - | - | - | + | + |
|  | *icsP* | - | + | + | + | - | - | + | + | + | + |
| Iron acquisition system | *iuc*C | + | + | + | + | + | + | + | + | + | + |
|  | *iut*A | + | + | + | + | + | + | + | + | + | + |
|  | *sit*A | + | + | + | + | + | + | + | + | + | + |
| Adhesion | *isc*A | - | + | + | + | - | + | - | - | + | + |
|  | *lpf*A | + | + | + | + | - | + | + | + | + | + |
|  | *pic* | - | - | - | - | - | - | - | - | - | - |
|  | *yeh*A | + | + | + | + | + | + | - | + | + | - |
|  | *yeh*B | + | + | + | + | + | + | + | + | + | + |
|  | *yeh*C | + | + | + | + | + | + | + | + | + | + |
|  | *yeh*D | + | + | + | + | + | + | + | + | + | + |
|  | *csg*A | + | + | + | + | + | + | + | + | + | + |

*Anr,* AraC negative regulator);*cap*U, Hexosyltransferase homolog); colE2, Colicin E2; colE7, Colicin E7; *csg*A, curlin major subunit CsgA; *gad*, Glutamate decarboxylase; *hly*E, Avian *E. coli*haemolysin; *ial*, invasion intestinal; *isc,* actin polymerizing factor; *inv*E, invasin E; *gtr*, glycosyltransferase; *iuc*C, Aerobactin synthetase; *iut*A, Ferric aerobactin receptor; *lpf*A, Long polar fimbriae; *nlp*I, lipoprotein NlpI precursor; *pic*, protein involved in intestinal colonization; *sen*B, Plasmid-encoded enterotoxin; sepA, serine protease adhesineA; *shi*A, homologs of the *Shigella flexneri* SHI-2 pathogenicity island gene shiA; *shi*B, homologs of the *Shigella flexneri* SHI-2 pathogenicity island gene *shi*A; *sig*A (serine protease autotransporters of Enterobacteriaceae (SPATE),*Shigella* IgA-like protease homologue; *sit*A, Iron transport protein; *stx*, shigatoxin; *ter*C, Tellurium ion resistance protein; *tra*T, Outer membrane protein complement resistance; *vir*A, VirA transmission from cell to another; *vir*B, VirB transcriptional activator; *vir*F, VirF transcriptional activator; *vir*K, VirK transcriptional activator; *yeh*A, Outer membrane lipoprotein, YHD fimbriael cluster; *yeh*B, Usher, YHD fimbriael cluster; *yeh*C, Chaperone, YHD fimbriael cluster; *yeh*D, Major pilin subunit, YHD fimbriael cluster; *ipa* operon, Invasion plasmid antigen; *osp* operon, Type III secretion system (T3SS) effectors; *ipg* operon, Type III secretion apparatus proteins; *ics* operon, biogenesis of cellular iron-sulfur protein-encoding genes; *Historic control strain from 2010
